# Nucleotide augmentation for machine learning-guided protein engineering

**DOI:** 10.1101/2022.03.08.483422

**Authors:** Mason Minot, Sai T. Reddy

## Abstract

Machine learning-guided protein engineering is a rapidly advancing field. Despite major experimental and computational advances however, collecting protein genotype (sequence) and phenotype (function) data remains time and resource intensive. As a result, the quality and quantity of training data is often a limiting factor in developing machine learning models. Data augmentation techniques have been successfully applied to the fields of computer vision and natural language processing, however, there is a lack of such augmentation techniques for biological sequence data. Towards this end we develop nucleotide augmentation (NTA), which leverages natural nucleotide codon degeneracy to augment protein sequence data in a biologically meaningful way. As a proof of concept for protein engineering, we apply NTA to train machine learning models with benchmark data sets of protein genotype and phenotype, revealing performance gains on par and surpassing benchmarks models, even when only using a fraction of the training data. NTA also enables substantial improvements for classification tasks under heavy class imbalance.

**Availability and implementation:** The code to use NTA and to reproduce the analyses in this study is publicly available at https://github.com/minotm/NTA

## Introduction

The application of machine learning (ML) on biological sequence data has expanded substantially in recent years^1,2^. One area of interest is ML-guided protein engineering, which enables efficient and large-scale exploration of protein sequence space^3^. This approach has been used for a variety of applications such as increasing protein expression^4^, multi-parameter optimization of antibody therapeutics^5^, improving the thermostability and function of enzymes^6,7^.

ML-guided protein engineering has been supported in recent years by advances in DNA synthesis, high-throughput screening assays and deep sequencing, which enable the generation of phentoype-genotype training data. However, collecting protein sequence and function (labeled) data still remains time- and resource-intensive. And as is the case in other applications of ML, small or imbalanced training data sets can often lead to bias and overfitting, thereby leading to a lack of performance and generalizability^8^. Moreover, large-scale data sets are often required to train effective deep learning models ^9,10^. Several approaches have been developed to make the most of limited data. For example, Wittman et al. (2021) show that ML-informed design of training data sets can improve directed evolution workflows^11^. Additionally, language models trained on large-scale and mostly natural sequence data have been developed to generate protein embeddings capable of improving downstream tasks like secondary structure and contact map prediction^7,12–15^.

In the ML-related fields of computer vision^8,16,17^ and natural language processing (NLP)^18–20^, data augmentation is commonly applied to combat data limitations. Data augmentation refers to techniques that artificially increase the number of training examples, which can lead to improved performance and act as a regularizer in reducing overfitting. Common image augmentation approaches include copying and warping an image, i.e via cropping and rotation^8^. NLP augmentation techniques may include copying a sentence and substituting words with synonyms to preserve meaning or translating a sentence into another language and back again^18–20^. Additionally synthetic data can be generated through a variety of techniques including Generative Adversarial Networks (GANs) and the Synthetic Minority Oversampling Technique (SMOTE)^8,21^.

Recently, data augmentation techniques have been developed for protein sequence data as well. Such approaches include GANs^22,23^, and augmentation for protein language models with contrastive learning via evolutionary information and string manipulations such as amino acid replacement and sequence shuffling^24,25^. While certainly noteworthy, these approaches are constrained to the domain in which they were trained (GANs), or may be less relevant in protein engineering workflows that generate large libraries of non-natural mutations in a small number of residues of a single protein^7,26,27^ (language models). Unlike computer vision and NLP, there exists a lack of simple, easy-to-apply data augmentation techniques for protein sequence data; likely resulting from the relationship between the discrete amino acid sequence of a protein and its structure and function.

Here we establish Nucleotide Augmentation (NTA), which represents a rapid and facile data augmentation technique for ML-guided protein engineering. By taking advantage of natural codon degeneracy (trinucleotide combinations), we developed NTA as the reverse translation of an amino acid sequence into multiple, unique nucleotide sequences via codon degeneracy (Fig. 1 and Algorithm 1). To benchmark and validate the performance of NTA, we select three protein engineering data sets with various sizes of training data and class balances^5,26,28,29^. Using benchmarked train/test splits, we determine that NTA can improve predictive performance of ML models with limited training data and under high class imbalance.

**Fig 1.**
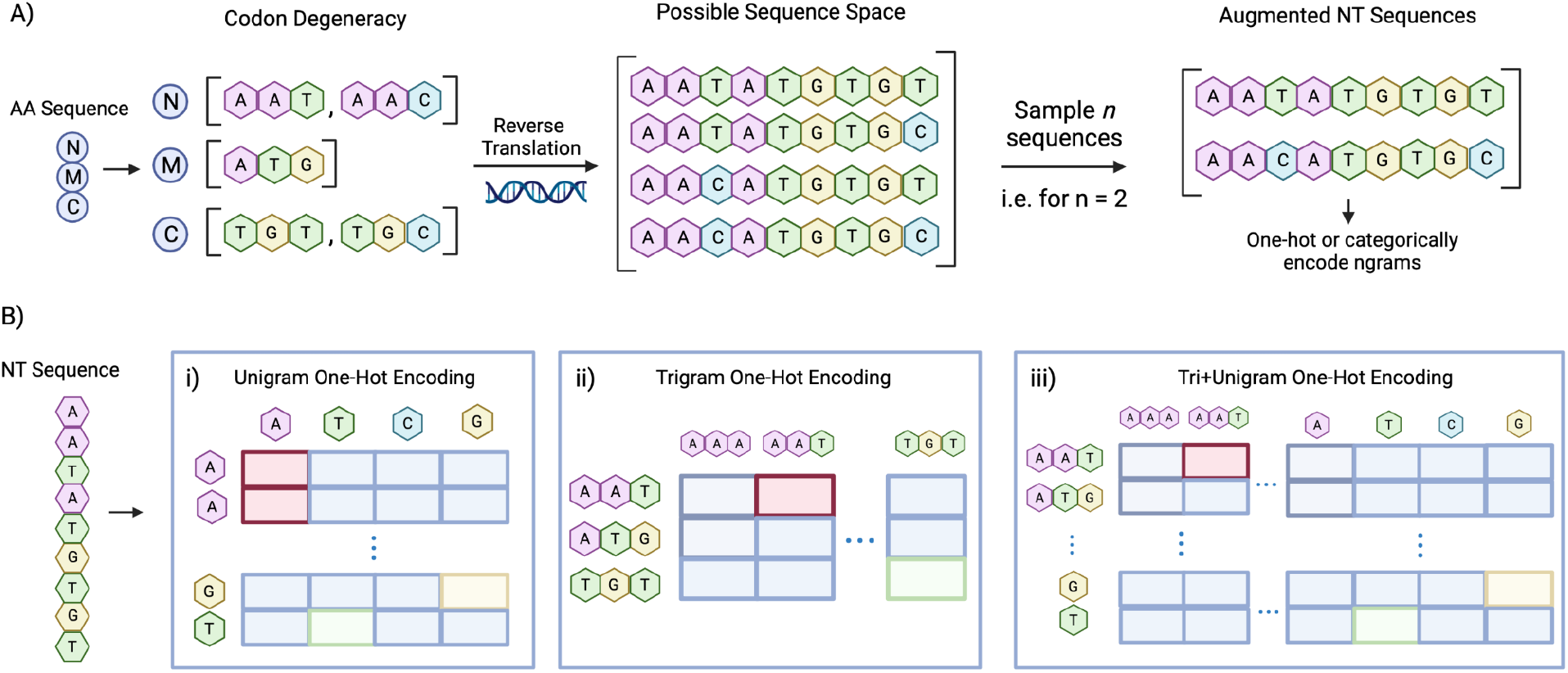
Nucleotide Augmentation Procedure and Encodings Tested. **A)** Nucleotide Augmentation (NTA) approach. Possible nucleotide codons are determined for each residue in an input amino acid sequence. Full-length nucleotide sequences are then sampled *n* times from the possible sequence space, where *n* is a user-specified augmentation factor. The sequences are then either categorically encoded (transformer) or one-hot encoded (CNN). The different ngram one-hot encoding approaches are illustrated in **B)**. First, a nucleotide sequence is broken into **B_i_)** unigrams **B_ii_)** trigrams or **B_iii_)** trigrams concatenated with unigrams (tri+unigrams). The ngrams are then tokenized according to their respective vocabularies. The tokenized vector corresponds to the transformer model’s input, however, this vector undergoes an additional one-hot encoding step for the CNN. For example, the resulting one-hot matrix size for a nucleotide sequence of length 9 will either be 9×4 (unigrams **B_i_**), 3×64 (trigrams, **B_ii_**), or 12×68 (tri+unigrams **B_iii_**). Note that amino acids are encoded as unigrams i.e. for sequence length L the resulting one-hot matrix is L×20 for the 20 canonical amino acids.

### Algorithm 1.

Nucleotide Augmentation Algorithm

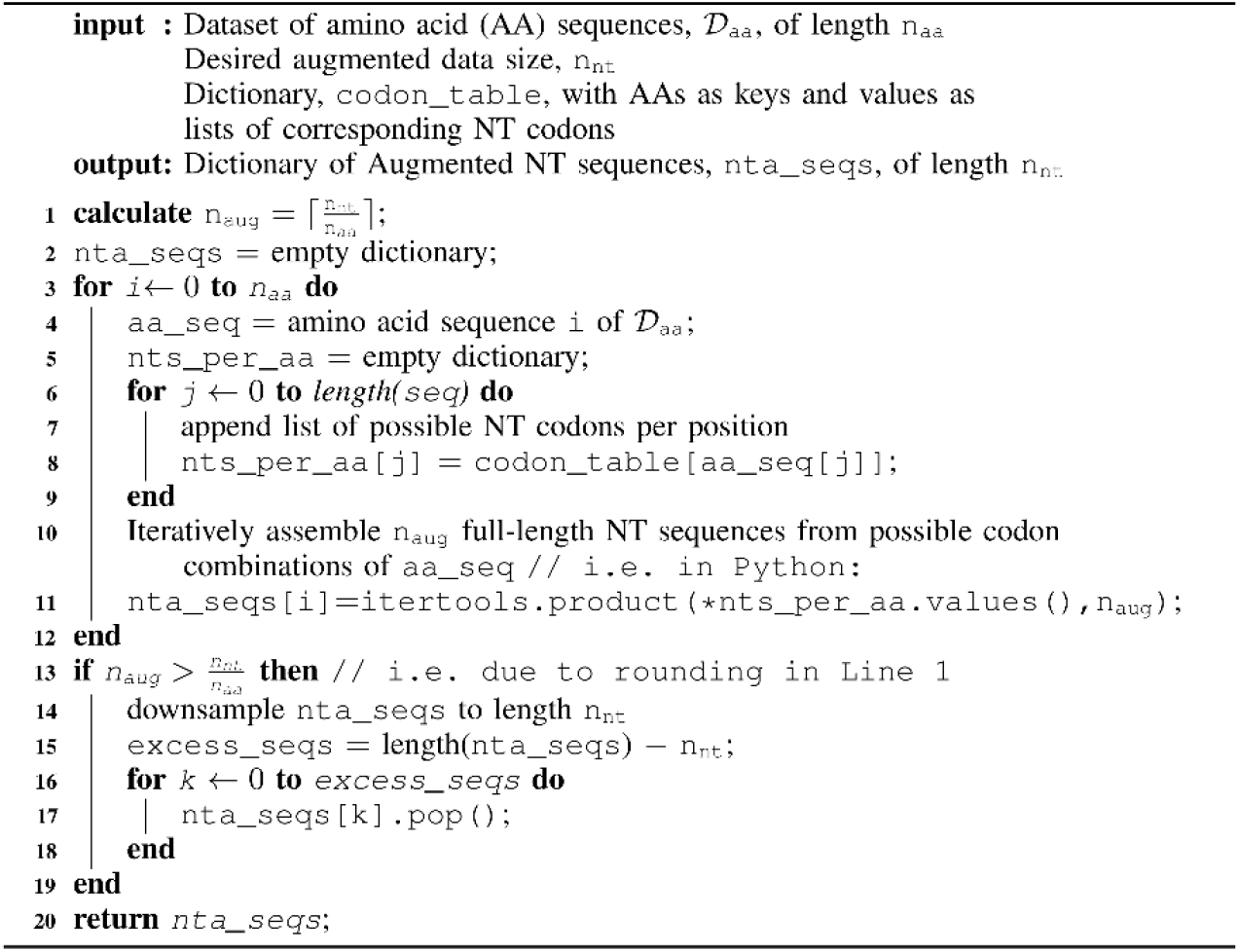

## Materials and Methods

### Data Sets Used In This Study

In order to evaluate the performance of NTA for ML-guided protein engineering, we acquired three benchmark labeled data sets. Such data sets consist of protein mutagenesis libraries where each sequence variant (genotype) in the library and its corresponding function (phenotype) are known. High-throughput screening by directed evolution coupled to deep sequencing provides an exemplary approach to generate such data sets. Although protein engineering has largely lacked benchmark data sets for ML, Dallego and colleagues recently published the Fitness Landscape Inference for Proteins (FLIP) repository^26^, which seeks to address this issue. We make use of two labeled regression data sets from FLIP as well as a third classification data set from a study related to ML-guided antibody engineering^5^ (Table 1).

**Table 1.** Description of data sets used in this study. Data Set corresponds to the name of the wild type (WT) protein that served as a basis for mutagenesis library generation. The edit distance threshold used to split the training/validation and test sets as well as the respective training, validation, and test data set sizes are included. *Length of AAV mutagenized region is listed as variable, as this library contains both insertions and deletions.

| Data Set | Source | Edit Distance from WT for Train/Test Split | Length of mutagenized region | # Training Set Sequences | # Validation Set Sequences | # Test Set Sequences |
| --- | --- | --- | --- | --- | --- | --- |
| GB1 | Wu et al. 2016 | 3 | 4 | 2,691 | 299 | 5,743 |
| AAV | Bryant et al. 2021 | 7 | variable*<br>(27 - 42) | 62,631 | 7,001 | 12,581 |
| Trastuzumab | Mason et al. 2021 | 7 | 9 | 11,172 | 2,795 | 3,858 |

#### GB1 Data Set

In the original study from Wu et al. (2016)^28^, four positions of GB1, the IgG-Fc binding domain of Protein G, were subjected to saturation combinatorial mutagenesis, thus resulting in an overall library diversity of 20^4^ = 160,000 variants. mRNA display, deep sequencing, and statistical analysis were used to determine fitness of the variants. The FLIP subtask chosen for this data set was ‘Three vs Rest’, a regression task that seeks to predict a value for variant fitness by training (and validating) on sequences with amino acid edit distance 1-3 (ED_1-3_) away from WT (wild type) and testing on sequences ED_4_ from WT. The subtask was chosen as the quantity of training data is large enough to allow investigation of how training sets of varying size impact model performance when supplemented with NTA. The full data set for this subtask includes 2,691 training, 299 validation, and 5,743 test sequences. Protein sequences are truncated to the variable region only.

#### AAV Data Set

Bryant et al. (2021)^29^ performed mutagenesis on a 29 amino acid region of the AAV capsid protein. Some variants contain insertions or deletions, resulting in a maximum of 39 mutations from WT. Variant fitness was assessed via viral production assay, deep sequencing and statistical analysis. The FLIP regression subtask chosen for this data set was ‘Seven vs Rest’ and seeks to predict a value for variant fitness. This subtask uses variants with an ED_1-7_ from WT as training and validation data. The test data set includes variants with ED_8-39_ from WT. The full data set for this subtask consists of 62,631 training, 7,001 validation and 12,581 test sequences. Protein sequences are truncated to the variable region only. This data set was chosen to complement the GB1 data, as it offers a larger, more complex training set with a higher number of mutated residues of variable length.

#### Trastuzumab Data Set

Mason et al. (2021)^5^ performed mutagenesis on 9 residues of the heavy chain complementarity determining region 3 (CDRH3) of the therapeutic antibody trastuzumab. Variants were screened for binding or non-binding to the HER2 antigen via mammalian display, fluorescence-activated cell sorting and deep sequencing. A train/test splitting strategy was developed specifically for this study. To correspond with the other data sets and to maximize the data used, ED_7_ from WT was chosen as the cutoff between train/validation and test sets. Using edit distance to split training and testing also resembles real-world workflows, in which models are trained with a limited number of mutations and used to extrapolate to a larger sequence space. The resulting training set was then balanced by downsampling the number of negatives (non-binders) to match the number of positives (antigen-binders) to create a balanced starting point for the synthetic introduction of class imbalance. 20% of the sequences were allocated as a validation set, resulting in a training set of 11,172 sequences, a validation set of 2,795 sequences, and a test set containing a balanced 1,929 positive and 1,929 negative sequences. Protein sequences are truncated to the variable region only.

### Nucleotide Augmentation Algorithm

For speed and reproducibility purposes, we performed NTA offline, prior to training, thus generating separate csv files for each condition. Psuedocode describing the NTA procedure is provided in Algorithm 1. In short, a user begins by specifying the augmentation factor, n_aug_, or number of augmentations to perform for each protein sequence. Next, all possible nucleotide codons for each amino acid in a sequence are determined and assembled into an ordered list of lists (or dictionary). n_aug_ is rounded up to the nearest integer and full-length nucleotide sequences are sampled n_aug_ times. This process is repeated for each protein sequence in the data set. In cases where n_aug_ is a float prior to rounding, down sampling is executed by removing one nucleotide sequence per unique amino acid sequence until the desired data set size is achieved. It should also be noted that augmentation capacity is limited by the diversity of the corresponding nucleotide codons. When given an n_aug_ that exceeds this upper bound, the algorithm returns the maximum number of unique sequences possible.

### Machine Learning Models

A transformer^30^ and a convolutional neural network (CNN) were chosen as representative models for this study due to their widespread use in ML on biological sequence data^5,12,13,26^. As the purpose of this work is not to develop the best model for the tasks, but to illustrate the potential of NTA, minimal hyperparameter optimization was performed and the same hyperparameters for each model are used across tasks. It is therefore reasonable to expect additional performance improvement with optimized parameters. The models were written in python using the PyTorch^31^ framework.

#### Transformer Model

A modified transformer^30^ was created with the following architecture. Protein and nucleotide sequences broken into ngrams are categorically encoded as input to an embedding layer with dimension 32 followed by positional encoding injection^30^ and connected to a transformer encoder layer with 2 attention heads, each with hidden dimension 128. The encoder output is first flattened, then fed to a linear layer with output dimension 512, followed finally by a linear layer with output dimension 1 to predict variant fitness or class. Rectified Linear Unit (ReLU) is applied as the activation function and a dropout of 0.3 are used throughout the network. To account for variable sequence length resulting from insertions and deletions in the AAV data, sequences are padded to the maximum length and a padding mask is used to prevent the attention mechanism from attending to padded tokens.

#### CNN Model

The CNN architecture is adapted and slightly scaled down from what is described in the FLIP repository model, which was applied to much larger data sets^26^. For the modified CNN, protein and nucleotide sequences broken into ngrams are one-hot encoded and used as input. The network consists of a 1D convolution with a kernel width of 5 and 64 convolutional filters, batch normalization and max pooling, mapping to a linear layer with 512 nodes and a dropout of 0.3, and final mapping to a linear layer with output dimension 1. ReLU is applied as the activation function throughout the network. Similar to the transformer, sequence padding and mask are applied to the AAV data set to handle sequences of variable length.

#### ngram Sequence Encoding

NTA requires reverse translation of amino acids to nucleotides, which increases the sequence length threefold. To probe how input format affects performance, three nucleotide ngram encoding schemes were tested: unigrams, trigrams, and tri+unigrams, which concatenates trigrams and unigrams. Each encoding scheme has a different vocabulary, or set of unique ‘words’ (ngrams). The unigram vocabulary has 4 elements: A,T,G,C. The trigram vocabulary consists of the 64 possible nucleotide codons. The tri+unigram vocabulary combines the two for a total of 68 elements. Protein sequences were encoded as unigrams with a vocabulary of 20, one character for each of the canonical amino acids. It is worth noting that the nucleotide trigram and amino acid unigram encodings share the same sequence length, albeit a different vocabulary size. After breaking a sequence into ngrams, categorical encoding and one-hot encoding are applied for the transformer and CNN, respectively.

#### Model Training

The loss function used for the regression tasks is Mean Squared Error and in accordance with previous work with the FLIP repository^26^, Spearman’s Rho is used to assess model performance. Binary Cross Entropy is used for the classification task and the Matthews Correlation Coefficient (MCC), which ranges from −1 to 1, is selected as the performance metric as it is an appropriate summary metric for binary classification of imbalanced data sets^32^. Stochastic gradient descent (SGD) with momentum of 0.9 is used as the optimizer. Training proceeds with a maximum of 250 epochs with 25 epochs early stopping patience for the GB1 and AAV tasks. Training for the antibody task proceeds 400 epochs with 40 epochs early stopping patience. A minimum of 40 epochs must be run prior to early stopping for all tasks. Mini-batch sizes 32 and 256 are used for the transformer and CNN, respectively. Models were trained using the ETH Zurich Euler Cluster with 1 GPU (Nvidia GTX 1080, GTX 1080 Ti, or V100) and a requested 16GB memory.

## Results

### Regression Variant Fitness Prediction With NTA

To probe how data quantity impacts ML models, it is common to truncate a training set to differing sizes and assess model performance. This type of analysis is useful since biological data collection is time and resource intensive, and thus can aid in experimental design.

To test how NTA performs with differing amounts of initial sequence diversity, training data sets were truncated into separate subsets consisting of 1%, 5%, 10%, 25%, 50%, 75%, and 100% of the available training data. A 0.5% fraction of the AAV data set was also created due to the data set’s larger total size. The truncated sets correspond to a range of either 26 to 2,691 (GB1) or 313 to 62,631 (AAV) sequences. In an attempt to mirror the distribution found in the full training and validation sets (as defined by the FLIP repository), sequences were binned by fitness value, a continuous value approximating mutant fitness experimentally determined in each study. Truncation was performed using train_test_split with ‘stratify’ option from sklearn (version 0.23.2) to maintain the ratio of binned sequences during truncation.

Baselines were collected on the truncated and full-size non-augmented training sets in both amino acid and nucleotide formats. Each data set was then augmented to three different final sizes ranging from ten times its initial size to several hundred thousand sequences, essentially treating the extent of augmentation as a hyperparameter to be tuned. This upper bound was chosen as it is an order of magnitude greater than the largest data set. The exceptions to this were the 1% and 5% truncated GB1 data sets, in which available nucleotide sequence diversity was less than this amount, in which case the data was augmented as much as possible. Validation and test sets were reverse translated without applying augmentation, consistent with common practice. Finally, the trained models were used to predict fitness values on the complete test sets. Performance was assessed using Spearman’s Rho, corresponding to FLIP benchmark repository protocol, across 5 separate random seeds. The highest performers are selected from the three different augmented data sets for each initial training set size and plotted in Figures 2 and 3.

**Figure 2.**
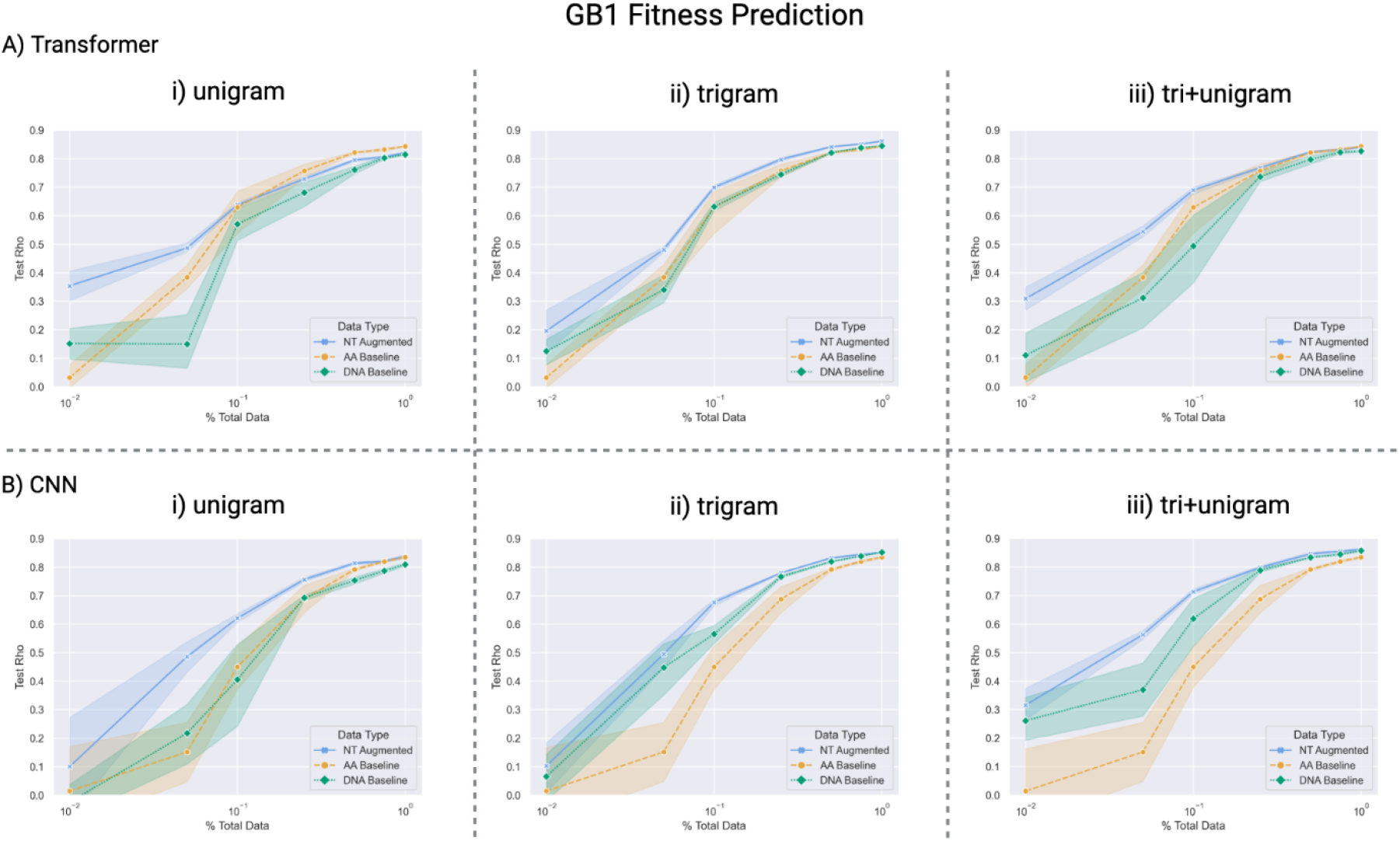
Impact of NTA on Regression Prediction of GB1 Fitness. Fitness prediction performance (Spearman’s Rho) of GB1 mutants as a function of the number of training samples used for model training. Points correspond to mean performance and shaded regions to 95 % confidence interval across 5 random seeds. Baselines include models trained on amino acid sequences (yellow) and models trained on DNA sequences without augmentation (green). Each fractionated data set was augmented to three different sizes, the best performing of which is reported as the blue X’s of the NT Augmented curve. **A)** Transformer model performance and **B)** CNN model performance when trained on unigrams (**A_i_**, **B_i_**), trigrams (**A_ii_**, **B_ii_**) and tri+unigrams (**A_iii_**, **B_iii_**).

**Figure 3.**
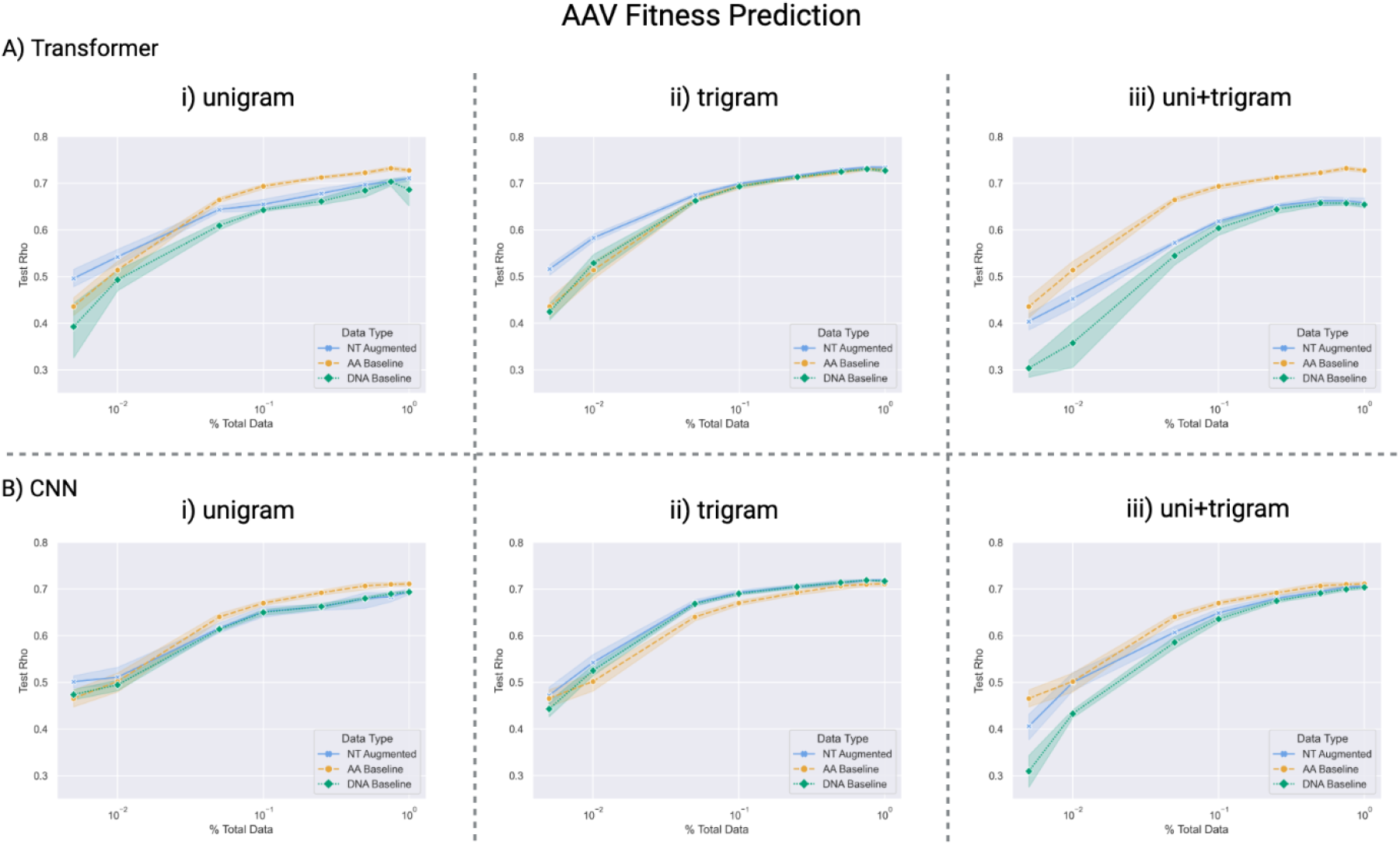
Impact of NTA on Regression Prediction of AAV Fitness. Fitness prediction performance (Spearman’s Rho) of AAV mutants as a function of the number of training samples used for model training. Points correspond to mean performance and shaded regions to 95 % confidence interval across 5 random seeds. Baselines include models trained on amino acid sequences (yellow) and models trained on DNA sequences without augmentation (green). Each fractionated data set was augmented to three different sizes, the best performing of which is reported as the blue X’s of the NT Augmented curve. **A)** Transformer model performance and **B)** CNN model performance when trained on unigrams (**A_i_**, **B_i_**), trigrams (**A_ii_**, **B_ii_**) and tri+unigrams (**A_iii_**, **B_iii_**).

**Figure 4.**
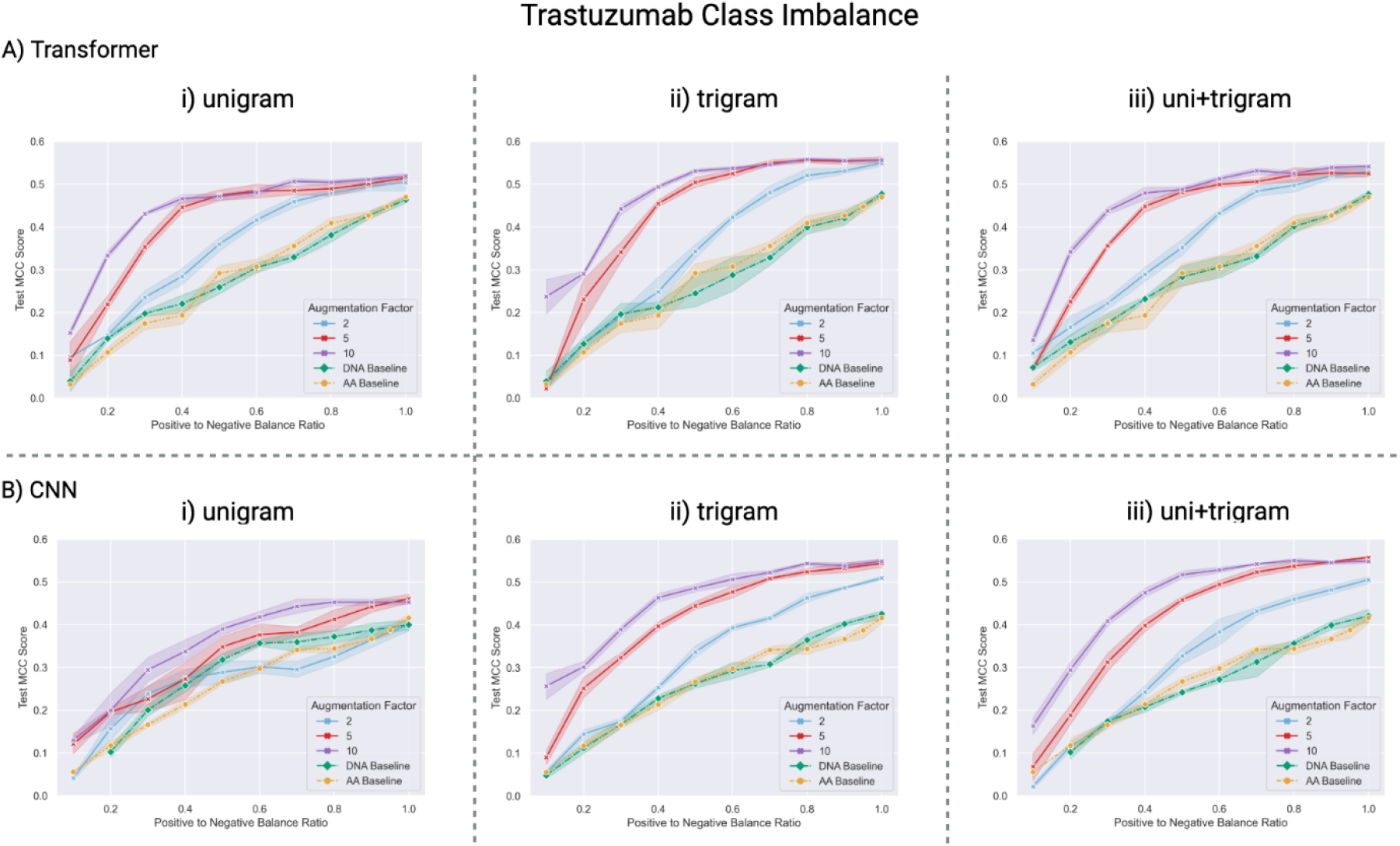
Imbalanced Classification of HER2 Binding of Trastuzumab Mutants. Binary classification (antigen binding and non-binding) prediction performance (Matthews Correlation Coefficient) of antibody variants as a function of the ratio of positive (minority class) to negative (majority class) sequences in the training set for the **A)** transformer and **B)** CNN models when trained on unigrams (**A_i_**, **B_i_**), trigrams (**A_ii_**, **B_ii_**) and tri+unigrams (**A_iii_**, **B_iii_**). NTA results in notable performance gains across models and encoding types. NTA is applied only to the minority (antigen binding) class and the majority class (antigen non-binding) is reverse translated from amino acids to nucleotides without augmentation. Augmentation factor refers to the number of unique nucleotide copies generated for every positive amino acid sequence in the training and validation sets. Performance on amino acid sequences (yellow) and non-augmented DNA sequences (green) are considered baselines.

### GB1 Mutant Fitness Prediction Results

NTA improves GB1 prediction for both the transformer and CNN models for almost every case, and notably for models trained with small data sets (Figure 2). For example, the unigram transformer on 1% of training data (25 sequences) results in a poor test Rho of 0.03 ± 0.05 on the AA baseline, and 0.13 ± 0.10 on the DNA baseline, however, NTA is able to improve performance to 0.35 ± 0.06 (Fig 2A_i_). Similarly, the tri+unigram CNN with 5% training data (134 sequences) results in a test Rho of 0.35 ± 0.18 on the DNA baseline, 0.15 ± 0.13 on the AA baseline, and 0.56 ± 0.02 using NTA (Figure 2B_iii_). It is also worth noting that NTA continues to result in improvements for most scenarios using larger training sets. The best performing model described in Dallego et al. (CNN) achieves 0.83 on the full FLIP data set. Remarkably, using only 50% training data and NTA, our CNN achieves 0.81 ± 0.00, 0.83 ± 0.00, 0.85 ± 0.00, with unigram, trigrams, and tri+unigrams, respectively.

### AAV Variant Fitness Prediction Results

ML models using NTA on the AAV data set generally result in notable improvements over the DNA baseline, the magnitude of which decreases as the number of initial samples is increased. For example, trigram transformer trained using NTA on .05% data (314 sequences) results in a Rho value of 0.51 ± 0.01, which is significantly higher than 0.44 ± 0.03 and 0.37 ± 0.08 for the AA and DNA baselines, respectively (Fig 3A_ii_). In general, NTA results in improvements over the DNA baseline, however, the DNA baseline is impacted by encoding choice to a greater extent than was observed on the GB1 data set. For unigrams and tri+unigrams, the DNA baseline is generally lower than training via amino acids. It is worth noting that both augmented and non-augmented DNA trigrams perform on par or better than the amino acid baseline. The best performing benchmark from Dallego et al^26^ is a CNN trained on the full FLIP data set with a Rho value of 0.74 . Although we performed virtually no hyperparameter optimization, our transformer trigram model trained with NTA on only 75% of available data is able to achieve a similar Rho of 0.73 ± 0.00.

### Class-Imbalanced Binding Classification With NTA

Protein engineering through the screening of mutagenesis libraries often results in a substantial fraction of low- or non-functional variants and a comparatively low fraction of variants with enhanced properties^11^. To test if NTA can improve learning on data with heavy class imbalance, we first create a balanced training set (Trastuzumab Data Set section) then artificially imbalance the data by down-sampling the number of positive examples (antibody variants binding to target antigen) to range from 10% to 100% of the number of negative sequences (antibody variants non-binding to antigen). Similar to our regression experiments, following downsampling, validation sets were split using sklearn’s ‘train_test_split’ function with ‘stratify’ option to produce validation sets 20% the size of each training set while maintaining the respective class imbalance. Baselines were collected on the imbalanced data for both amino acid and non-augmented DNA sequences. NTA was applied only to the minority class (antigen-binding sequences) of the training set in accordance with common practice^33–35^. The minority class was augmented by a factor of 2, 5, and 10. Further augmentation beyond 10 generally caused a drop in performance. The majority class (antigen non-binding) sequences were reverse translated without augmentation. PyTorch’s WeightedRandomSampler was used to class balance mini-batches as much as possible during training as this was found to result in better performance.

### Imbalanced Classification Results

NTA yields significant improvements on imbalanced antibody-antigen binding data for every model-encoding scheme tested. In general, the baseline and augmented transformer models outperform the CNN. NTA results in significant gains under heavy class imbalance. For example with a 0.3 positive to negative ratio, NTA improves MCC from 0.17 ± 0.03 and 0.19 ± 0.02 (AA and DNA Baselines, respectively) to 0.44 ± 0.02 at an augmentation factor of 10 with the trigram transformer (Fig 3A_ii_). Performance generally increases with an augmentation factor up to 10, at which point performance was seen to drop as augmentation was pushed further (data not shown). The trigram transformer was found to perform the best and the unigram CNN the poorest. Overall, these results demonstrate the ability of NTA to aid in learning from class-imbalanced data.

## Discussion

The collection of protein genotype-phenotype data is time and resource intensive, thus representing a critical bottleneck for ML-guided protein engineering. Previous work has sought to counter data limitations with GANs^22,23^ and language models^24,25^, however, GANs are limited to the domain in which they are trained and language models may not always be appropriate for protein engineering workflows introducing large libraries of non-natural mutations into a small number of residues of a protein^7,26,27^. For example, Dallago and colleagues (2021)^26^ find that for synthetic mutagenesis data sets focusing on a single protein, language models can be outperformed by smaller, more focused models. To date the field lacks the simple, easy-to-use data augmentation techniques that are commonly found in the fields of computer vision and NLP. Towards this end, we develop here NTA, which leverages the natural nucleotide codon degeneracy of protein sequences to augment data sets for ML. As a proof of concept, we apply NTA to three labeled data sets for ML-guided protein engineering, two of which (GB1 and AAV) have been established as benchmarks by the recent FLIP repository^26^.

We find that NTA yields large gains when limited training data is available (e.g., 10 − 10^3^ sequences), while still enhancing performance as more data is collected. This is particularly useful for protein engineering workflows unable to generate high throughput data (i.e. variant stability and affinity characterization or cell-based assays). We also find that NTA serves as a promising method to improve learning on class imbalanced data, which is a common occurrence in protein engineering experiments^11^.

NTA advantages include that it is biologically meaningful, can be easily applied to most protein-ML workflows, and requires minimal additional resources. One potential downside of NTA is that it prohibits the use of models pretrained with amino acids. It is worth noting that NTA could be applied to additional areas of ML-guided protein engineering, such as predictions of structure, stability, immunogenicity and subcellular localization^13,36^. Finally, NTA could supplement contrastive learning and be combined with or used to aid in the training of protein language models^24,25^. Future work may also seek to improve the NTA algorithm and its application for specific use cases.

## Acknowledgements

We thank Margarita Pertseva and Anna Weber for valuable scientific discussions.

Figures were created using BioRender.com.

## References

1. Jurtz VI, Johansen AR, Nielsen M, et al. An introduction to deep learning on biological sequence data: examples and solutions. Bioinformatics. 2017;33(22):3685–3690. doi:10.1093/bioinformatics/btx531

2. Deep learning for computational biology. Mol Syst Biol. 2016;12(7):878. doi:10.15252/msb.20156651

3. Yang KK, Wu Z, Arnold FH. Machine-learning-guided directed evolution for protein engineering. Nat Methods. 2019;16(8):687–694. doi:10.1038/s41592-019-0496-6

4. Shin JE, Riesselman AJ, Kollasch AW, et al. Protein design and variant prediction using autoregressive generative models. Nat Commun. 2021;12(1):2403. doi:10.1038/s41467-021-22732-w

5. Mason DM, Friedensohn S, Weber CR, et al. Optimization of therapeutic antibodies by predicting antigen specificity from antibody sequence via deep learning. Nat Biomed Eng. 2021;5(6):600–612. doi:10.1038/s41551-021-00699-9

6. Romero PA, Krause A, Arnold FH. Navigating the protein fitness landscape with Gaussian processes. Proc Natl Acad Sci. 2013;110(3):E193–E201. doi:10.1073/pnas.1215251110

7. Biswas S, Khimulya G, Alley EC, Esvelt KM, Church GM. Low-N protein engineering with data-efficient deep learning. Nat Methods. 2021;18(4):389–396. doi:10.1038/s41592-021-01100-y

8. Shorten C, Khoshgoftaar TM. A survey on Image Data Augmentation for Deep Learning. J Big Data. 2019;6(1):60. doi:10.1186/s40537-019-0197-0

9. The Unreasonable Effectiveness of Data. Accessed December 12, 2021. https://www.computer.org/csdl/magazine/ex/2009/02/mex2009020008/13rRUy0HYOb

10. Sun C, Shrivastava A, Singh S, Gupta A. Revisiting Unreasonable Effectiveness of Data in Deep Learning Era. In: 2017 IEEE International Conference on Computer Vision (ICCV). ; 2017:843–852. doi:10.1109/ICCV.2017.97

11. Wittmann BJ, Yue Y, Arnold FH. Informed training set design enables efficient machine learning-assisted directed protein evolution. Cell Syst. 2021;12(11):1026–1045.e7. doi:10.1016/j.cels.2021.07.008

12. Rao RM, Liu J, Verkuil R, et al. MSA Transformer. In: Proceedings of the 38th International Conference on Machine Learning. PMLR; 2021:8844–8856. Accessed March 7, 2022. https://proceedings.mlr.press/v139/rao21a.html

13. Rao R, Bhattacharya N, Thomas N, et al. Evaluating Protein Transfer Learning with TAPE. In: Wallach H, Larochelle H, Beygelzimer A, Alché-Buc F d\textquotesingle, Fox E, Garnett R, eds. Advances in Neural Information Processing Systems 32. Curran Associates, Inc.; 2019:9689–9701. Accessed June 20, 2020. http://papers.nips.cc/paper/9163-evaluating-protein-transfer-learning-with-tape.pdf

14. Luo Y, Jiang G, Yu T, et al. ECNet is an evolutionary context-integrated deep learning framework for protein engineering. Nat Commun. 2021;12(1):5743. doi:10.1038/s41467-021-25976-8

15. Ofer D, Brandes N, Linial M. The language of proteins: NLP, machine learning & protein sequences. Comput Struct Biotechnol J. 2021;19:1750–1758. doi:10.1016/j.csbj.2021.03.022

16. Perez L, Wang J. The Effectiveness of Data Augmentation in Image Classification using Deep Learning. ArXiv171204621 Cs. Published online December 13, 2017. Accessed November 23, 2021. http://arxiv.org/abs/1712.04621

17. Taylor L, Nitschke G. Improving Deep Learning with Generic Data Augmentation. In: 2018 IEEE Symposium Series on Computational Intelligence (SSCI). ; 2018:1542–1547. doi:10.1109/SSCI.2018.8628742

18. Feng SY, Gangal V, Wei J, et al. A Survey of Data Augmentation Approaches for NLP. In: Findings of the Association for Computational Linguistics: ACL-IJCNLP 2021. Association for Computational Linguistics; 2021:968–988. doi:10.18653/v1/2021.findings-acl.84

19. Sennrich R, Haddow B, Birch A. Improving Neural Machine Translation Models with Monolingual Data. In: Proceedings of the 54th Annual Meeting of the Association for Computational Linguistics (Volume 1: Long Papers). Association for Computational Linguistics; 2016:86–96. doi:10.18653/v1/P16-1009

20. Anaby-Tavor A, Carmeli B, Goldbraich E, et al. Do Not Have Enough Data? Deep Learning to the Rescue! Proc AAAI Conf Artif Intell. 2020;34(05):7383–7390. doi:10.1609/aaai.v34i05.6233

21. Chawla NV, Bowyer KW, Hall LO, Kegelmeyer WP. SMOTE: Synthetic Minority Over-sampling Technique. J Artif Intell Res. 2002;16:321–357. doi:10.1613/jair.953

22. Han X, Zhang L, Zhou K, Wang X. ProGAN: Protein solubility generative adversarial nets for data augmentation in DNN framework. Comput Chem Eng. 2019;131:106533. doi:10.1016/j.compchemeng.2019.106533

23. Li M, Zhang W. PHIAF: prediction of phage-host interactions with GAN-based data augmentation and sequence-based feature fusion. Brief Bioinform. 2021;(bbab348). doi:10.1093/bib/bbab348

24. Shen H, Price LC, Bahadori T, Seeger F. Improving Generalizability of Protein Sequence Models with Data Augmentations.; 2021:2021.02.18.431877. doi:10.1101/2021.02.18.431877

25. Lu AX, Lu AX, Moses A. Evolution Is All You Need: Phylogenetic Augmentation for Contrastive Learning. ArXiv201213475 Cs Q-Bio. Published online December 24, 2020. Accessed November 30, 2021. http://arxiv.org/abs/2012.13475

26. Dallago C, Mou J, Johnston KE, et al. FLIP: Benchmark Tasks in Fitness Landscape Inference for Proteins.; 2021:2021.11.09.467890. doi:10.1101/2021.11.09.467890

27. Wittmann BJ, Johnston KE, Wu Z, Arnold FH. Advances in machine learning for directed evolution. Curr Opin Struct Biol. 2021;69:11–18. doi:10.1016/j.sbi.2021.01.008

28. Wu NC, Dai L, Olson CA, Lloyd-Smith JO, Sun R. Adaptation in protein fitness landscapes is facilitated by indirect paths. Neher RA, ed. eLife. 2016;5:e16965. doi:10.7554/eLife.16965

29. Bryant DH, Bashir A, Sinai S, et al. Deep diversification of an AAV capsid protein by machine learning. Nat Biotechnol. 2021;39(6):691–696. doi:10.1038/s41587-020-00793-4

30. Vaswani A, Shazeer N, Parmar N, et al. Attention is All you Need. In: Advances in Neural Information Processing Systems. Vol 30. Curran Associates, Inc.; 2017. Accessed March 7, 2022. https://papers.nips.cc/paper/2017/hash/3f5ee243547dee91fbd053c1c4a845aa-Abstract.html

31. Paszke A, Gross S, Massa F, et al. PyTorch: An Imperative Style, High-Performance Deep Learning Library. In: Advances in Neural Information Processing Systems. Vol 32. Curran Associates, Inc.; 2019. Accessed December 12, 2021. https://papers.nips.cc/paper/2019/hash/bdbca288fee7f92f2bfa9f7012727740-Abstract.html

32. Chicco D, Jurman G. The advantages of the Matthews correlation coefficient (MCC) over F1 score and accuracy in binary classification evaluation. BMC Genomics. 2020;21(1):6. doi:10.1186/s12864-019-6413-7

33. Shamsolmoali P, Zareapoor M, Shen L, Sadka AH, Yang J. Imbalanced data learning by minority class augmentation using capsule adversarial networks. Neurocomputing. 2021;459:481–493. doi:10.1016/j.neucom.2020.01.119

34. Saini M, Susan S. Data Augmentation of Minority Class with Transfer Learning for Classification of Imbalanced Breast Cancer Dataset Using Inception-V3. In: Morales A, Fierrez J, Sánchez JS, Ribeiro B, eds. Pattern Recognition and Image Analysis. Lecture Notes in Computer Science. Springer International Publishing; 2019:409–420. doi:10.1007/978-3-030-31332-6_36

35. Afzal S, Maqsood M, Nazir F, et al. A Data Augmentation-Based Framework to Handle Class Imbalance Problem for Alzheimer’s Stage Detection. IEEE Access. 2019;7:115528–115539. doi:10.1109/ACCESS.2019.2932786

36. Li G, Iyer B, Prasath VBS, Ni Y, Salomonis N. DeepImmuno: deep learning-empowered prediction and generation of immunogenic peptides for T-cell immunity. Brief Bioinform. 2021;22(6):bbab160. doi:10.1093/bib/bbab160

